# Host range and genetic plasticity explain the co-existence of integrative and extrachromosomal mobile genetic elements

**DOI:** 10.1101/250266

**Authors:** Jean Cury, Pedro H. Oliveira, Fernando de la Cruz, Eduardo P.C. Rocha

## Abstract

Self-transmissible mobile genetic elements drive horizontal gene transfer between prokaryotes. Some of these elements integrate in the chromosome, whereas others replicate autonomously as plasmids. Recent works showed the existence of few differences, and occasional interconversion, between the two types of elements. Here, we enquired on why evolutionary processes have maintained the two types of mobile genetic elements by comparing integrative and conjugative elements (ICE) with extrachromosomal ones (conjugative plasmids) of the highly abundant MPF_T_ conjugative type. We observed that plasmids encode more replicases, partition systems, and antibiotic resistance genes, whereas ICEs encode more integrases and metabolism-associated genes. ICEs and plasmids have similar average sizes, but plasmids are much more variable, have more DNA repeats, and exchange genes more frequently. On the other hand, we found that ICEs are more frequently transferred between distant taxa. We propose a model where differential plasticity and transmissibility range explain the co-occurrence of integrative and extra-chromosomal elements in microbial populations. In particular, the conversion from ICE to plasmid allows ICE to be more plastic, while the conversion from plasmid to ICE allows the expansion of the element‘s host range.

## Introduction

The genomes of Prokaryotes have mobile genetic elements (MGEs) integrated in the chromosome or replicating as extrachromosomal elements. These MGEs usually encode non-essential but ecologically important traits (1, 2). Extra-chromosomal elements, such as conjugative plasmids (CPs) and lytic phages, replicate autonomously in the cell using specialized replicases to recruit the bacterial DNA replication machinery (or to use their own). Plasmids and extra-chromosomal prophages can also increase their stability in cellular lineages using partition systems, for proper segregation during bacterial replication (3), resolution systems, to prevent accumulation of multimers (4), and restriction-modification or toxin-antitoxin systems, for post-segregation killing of their hosts (5). Alternatively, many MGEs integrate into the chromosome. This is the case of the vast majority of known prophages, of most conjugative elements (ICEs), and of many elements with poorly characterized mechanisms of genetic mobility (*e.g.*, many pathogenicity islands)(6–8). The integrated elements are replicated along with the host chromosome and require an additional step of excision before being transferred between cells. The existence of both integrative and extra-chromosomal elements was a fruitful source of controversy in the dawn of molecular biology, eventually leading to the discovery of the molecular mechanisms allowing both states (9, 10). Yet, a complementary question does not seem to have been addressed in the literature: Why are there both types of elements? What are the relative benefits and disadvantages of the integrated and extrachromosomal MGE?

To address these questions, we analyzed the differences and similarities between ICEs and CPs. We focused on these elements because both forms are frequently found in bacteria, they can be easily detected in genomes, and the mechanism of conjugation is well known. Conjugative elements have a crucial role in spreading antibiotic resistance and virulence genes among bacterial pathogens (11–14). Recently, several works suggested that the line separating integrative ICEs and CPs could be thinner than anticipated (15), because some ICEs encode plasmid-associated functions like replication (16) or partition (17), some plasmids encode integrases (18), and ICEs and CPs are intermingled in the phylogenetic tree of conjugative systems (19). Finally, both forms – ICEs and CPs – are found throughout the bacterial kingdom, but their relative frequency depends on the taxa and on the mechanisms of conjugation (7). These differences suggest that conjugative elements endure diverse selective pressures for being integrative or extrachromosomal depending on unknown environmental, genetic, or physiological variables.

We thought that key differences in the biology of integrative and extrachromosomal elements might provide them with different types of advantages. ICEs require an additional step of integration/excision during transfer, which may take time and requires genetic regulation. Their integration in the chromosome may affect the latter‘s organization and structure, and these collateral effects might depend on the size of the element. On the other hand, ICEs replicate as part of chromosomes and could thus be lost from the cell at lower rates than plasmids. Furthermore, plasmids must recruit the host replication machinery, which may render elements incompatible and is known to constrain their host range: many plasmids are able to conjugate into distantly related hosts, but are unable to replicate there (20–22). We thus hypothesize that ICEs might be favored when transfers occur between distant hosts, whereas plasmids might provide more genetic plasticity because their size is not constrained by chromosomal organization.

Here, we study conjugative elements of the type MPF_T_. This is the most frequent and best-studied type of conjugative systems (19), and the only one for which we can identify hundreds of elements of each of the forms (ICEs and CPs). We restricted our analysis to genera containing both CPs and ICEs, to avoid, as much as possible, taxonomical biases. We first describe the content of both types of elements and highlight their differences and similarities. Next, we quantify their genetic similarity and the extent of their gene exchanges. Finally, we show that chromosomal integration facilitates the colonization of novel taxa by a conjugative element.

## Results

### Functional and genetic differences between ICEs and CPs

We analyzed a set of 151 ICEs and 136 CPs of the same genera and of type MPF_T_, most of which were from Proteobacteria (96.9%). Both ICEs and CPs were found to be AT-richer than their host chromosomes, which is a common feature in MGEs and horizontally transferred genes (23). However, the difference was three times smaller for ICEs (Fig. 1A), presumably because they replicate with the chromosome or remain a longer time in the same host. The average size of CPs is slightly larger (75kb vs 59kb), and the median slightly smaller (46kb vs 52kb) than that of ICEs. In contrast, CPs have more diverse sizes than ICEs (Fig. 1B), showing a coefficient of variation twice as large (1.05 vs 0.49). The size of the conjugative elements depends on the size of the bacterial genome (after discounting the size of their conjugative elements), this effect being much stronger for CPs (Fig. 1C). CPs also have four times higher density of large DNA repeats than ICEs (Fig. 1D). These results suggest that CPs diversify faster than ICEs.

**Fig. 1:**
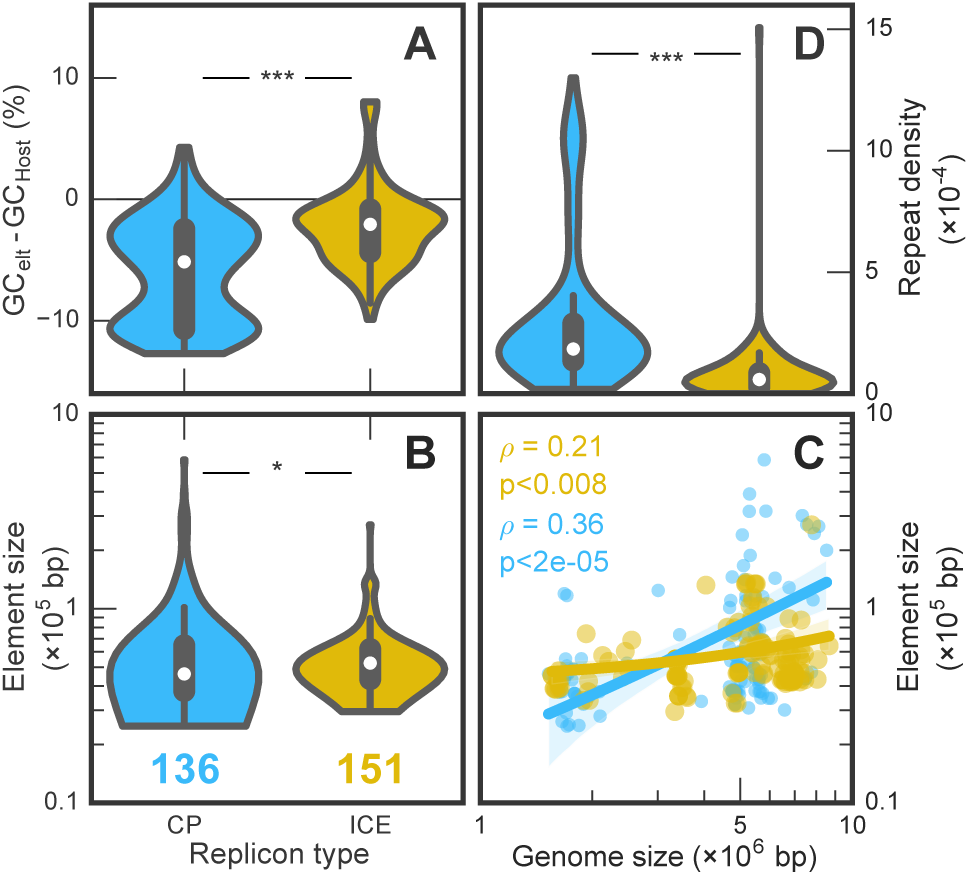
Comparison between 136 CPs (blue) and 151 ICEs (yellow) in composition and sizes. CPs are AT richer than ICEs relative to their hosts (Wilcoxon rank sum test, p-value<10^−3^).ICEs and CPs have different distributions of size (same test, p-value<0.05). Median sizes: ICEs (52.5 kb) > CPs (46.1 kb). Averages: ICEs (59 kb) < CPs (74.6 kb) **C.** Size of the element as a function of the genome size of its host (decreased by the size of the mobile element itself). Shaded regions indicate the 95% confidence interval. The Y-axis is identical to the one in panel B. **D**. Density of repeats is higher in CPs than in ICEs (0.30 vs 0.078 repeats per kb, same test, p-value < 10^−10^).

HGT concentrates in a few hotspots in bacterial chromosomes, presumably to minimize disruption in their organization (24). We used HTg50, a measure of the concentration of HGT in chromosomes that corresponds to the minimal number of spots required to account for 50% of horizontally transferred genes (24), to test if chromosomes with fewer integration hotspots had more plasmids. Indeed, there is a negative association between the number of plasmids, weighted by their size, and the chromosomes’ HTg50 (Spearman ρ=-0.35, p-value=0.0016, Fig. S1).

We then quantified the differences between ICEs and CPs in terms of functions associated with their biology, with a focus on stabilization functions. Relaxases are part of the rolling circle replication initiator proteins and some have been shown to act as replicases(16, 25) or site-specific recombinases (26, 27). Since all conjugative elements encode a relaxase, by definition, they may also have these functions. In the following, we focused on typical plasmid replication initiator proteins (more than 95% of them are involved in theta-replication, and none is matched by the protein profiles of relaxases), and serine or tyrosine recombinases as integrases. Expectedly, ICEs showed higher frequency of integrases, while CPs had more frequently identifiable partition and replication systems. Some ICEs encode partition systems (11%) and many encode a replicase (40%), while 37% of CPs encode at least one tyrosine or serine recombinase (Fig. 2A). These results further illustrate a continuum between the two types of elements: about half of the elements (40% and 48%, ICEs and CPs, respectively) have functions usually associated with the other type and may (rarely) lack functions typically associated with its own type (Fig. 2B). Interestingly, we observe that ICEs containing replication or partition systems contain more repeats per nucleotide than the others (Wilcoxon rank-sum test, p-value = 0.02). We identified plasmid incompatibility systems of diverse types, whereas ICE could not be typed in the current scheme (Fig. S2).

**Fig. 2:**
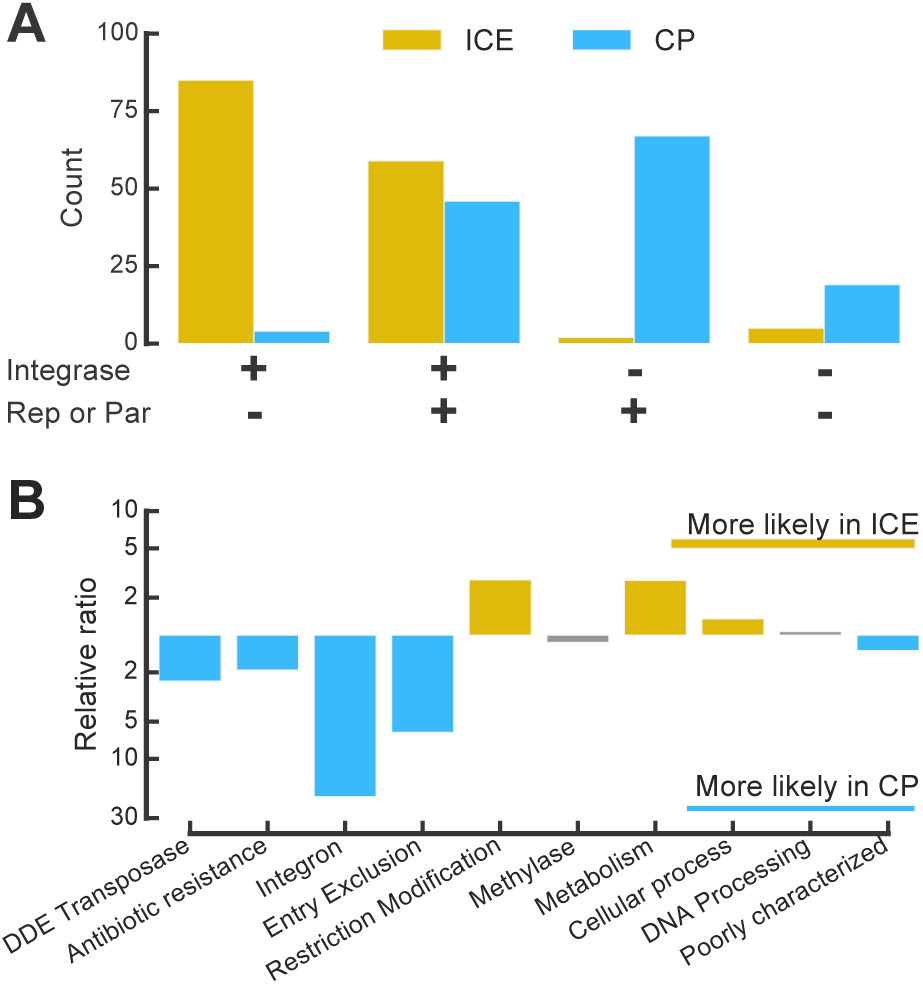
Comparisons of the functions carried by ICEs and CPs. **A.** Elements encoding (+) or lacking (-) replication or partition systems (“Rep or Par”) or an integrase. **B.** Accessory functions over-represented (yellow) or under-represented (blue) in ICEs relative to CPs. Colored bars: significantly different from zero, p-value < 0.05 Fisher exact test with Bonferroni-Holm correction for multiple tests. Grey bars otherwise.

We then made similar analyses for functions usually regarded as accessory or unrelated to the biology of MGEs (Fig. 2B). ICEs were more likely to carry restriction-modification systems (×2.8) than CPs (but not orphan methylases), suggesting that ICEs endure stronger selective pressure for stabilization in the genome. In contrast, they were significantly less likely to carry antibiotic resistance genes or integrons. They also had fewer identifiable entry-exclusion systems, which may reflect the ability of ICEs to tolerate the presence of multiple similar elements in the cell (28). The classification of genes in the four major functional categories of the EggNOG database, showed that ICEs had relatively more genes encoding metabolic and cellular processes. We have previously shown that genes of unknown or poorly characterized function were over-represented in ICEs relative to their host chromosome (29). The frequency of these genes is even higher in plasmids (61% vs 46%). Hence, both types of elements have many functions in common, but their relative frequency often differs significantly.

### Genetic similarities between ICEs and CPs

The results of the previous section, together with previously published studies (see Introduction), suggest that ICEs and CPs either share a common history or often exchange genes (or both). We detailed the relationships of homology between ICEs and CPs using the weighted Gene Repertoire Relatedness (wGRR), which measures the frequency of bi-directional best hits between two elements weighted by their sequence similarity (see Methods). We clustered the matrix of wGRR using the Louvain algorithm (30), and found six well-distinguished groups (Fig. 3). Two groups (1 and 6) are only constituted of CPs, two are composed of more than 90% of ICEs (3 and 5) and two have a mix of both types of elements (2 and 4) (Fig.3, top bar). Bacterial species are scattered between groups, showing that they are not the key determinant of the clustering. Some groups are only from γ-proteobacteria, but others include bacteria from different classes. Groups where elements are from the same taxonomic classes tend to have either CPs or ICEs, whereas the others have mixtures of both elements. Group 4, includes many ICEs and CPs, where all ICEs have integrases while more than half of the CPs lack both replication and partition systems (Fig. 4). This group includes almost only elements from ε-proteobacteria that may have specificities that we were not able to take into account. In contrast, almost all ICEs of groups 2 and 3 encode an integrase and all CPs have partition or replication systems.

**Fig. 3:**
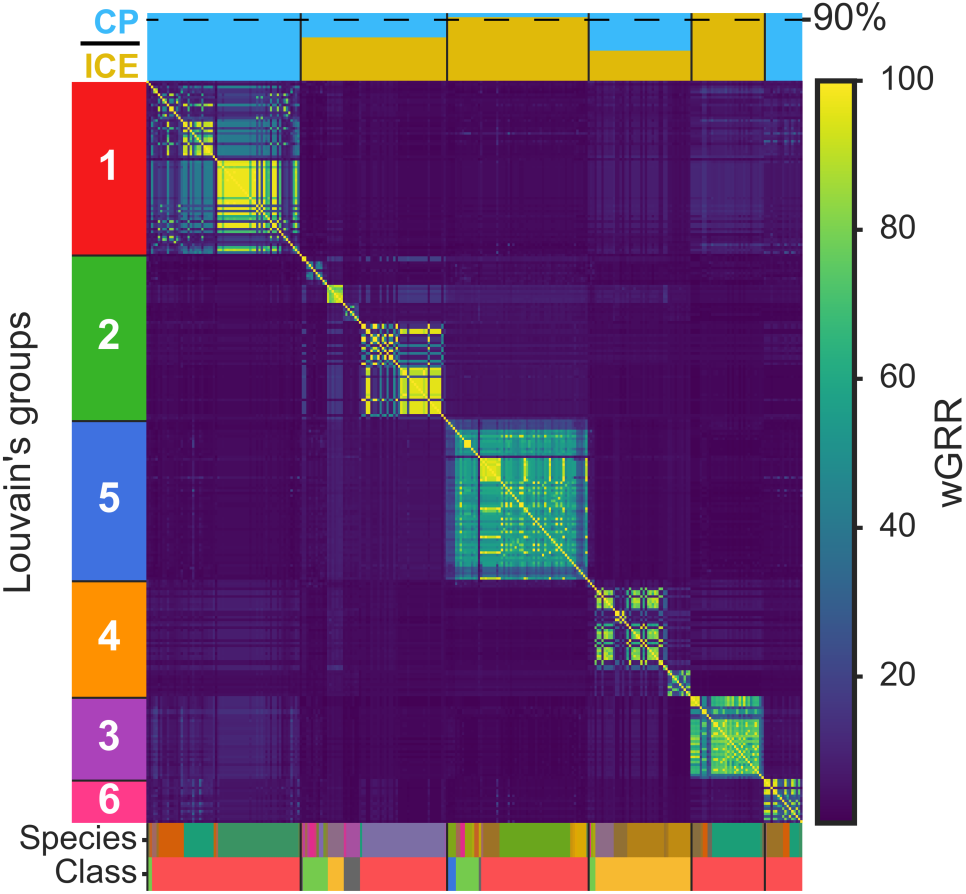
Heatmap of the wGRR scores, ordered after the size of Louvain‘s group (depicted on the left bar). The top bar represents the proportion of ICEs (yellow) and CPs (blue) for each group. The bottom bar assigns a color corresponding to the host‘s species or class (εproteobacteria in red, β-proteobacteria in green, α-proteobacteria in blue, ε- proteobacteria In orange, Fusobacteria and Acidobacteria in grey).

**Fig. 4:**
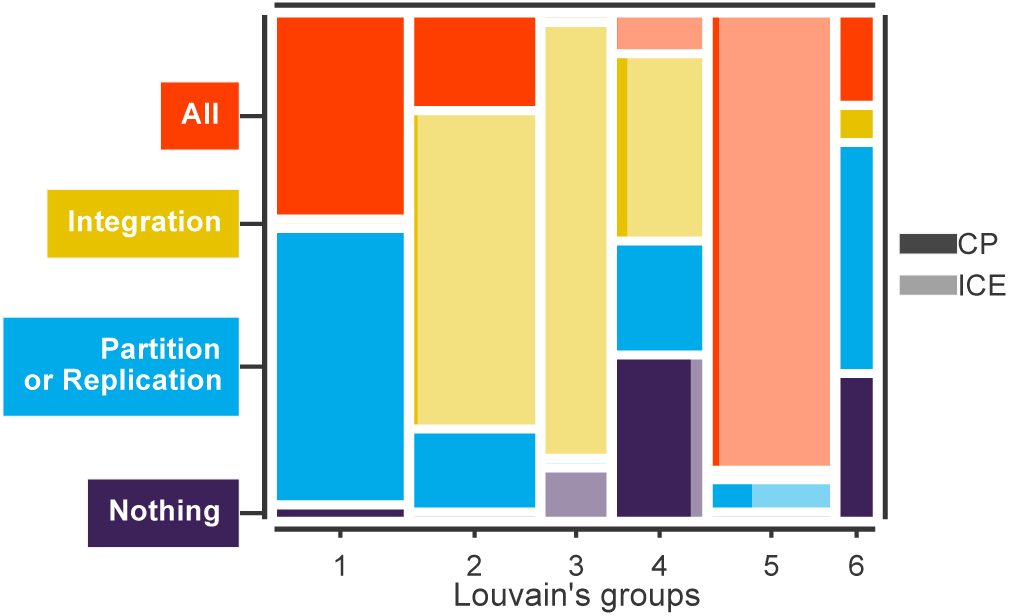
Mosaic plot representing the frequency of key functions of conjugative elements in terms of the Louvain‘s groups (Fig. 3). The width of the bar is proportional to the number of elements in a given Louvain‘s group (see the number of elements of each group on the top of the bars) and the areas reflect the proportion of the elements with the function. The colors represent the type of function, and their tint represent the part of ICEs (lighter) and CPs (darker).

We controlled for the effect of the MPF genes in the previous clustering analysis by re-doing it without these genes (Fig. S3). This produced the same number of groups - N1 to N6 - and 90% of the elements of the former groups were classed in the same novel groups (Fig. S4). The only qualitatively significant difference between the two analyses concerned the group 2 for which 36% of the elements were now classed in groups N4 or N6. Overall, these controls confirm that ICEs and CPs can be grouped together, and apart from other elements of the same type. The grouping is not caused by sequence similarity between conjugative systems. Instead, it probably reflects either within group genetic exchanges between ICEs and CPs, or interconversions of the two types of elements.

### Genetic exchanges: CPs become ICEs for broader host range

The clustering of ICEs and CPs could be explained by genetic transfer between them. To address this question, we searched for pairs of conjugative elements with low wGRR (<30%) but some highly similar homologs (>80% sequence identity). This identified several cases of recent transfer of a single or a few genes between elements (Fig. 5A). In agreement with our observations of higher genetic plasticity in CPs, most transfers took place between these types of elements (Fig. 5.B-D).

**Fig. 5:**
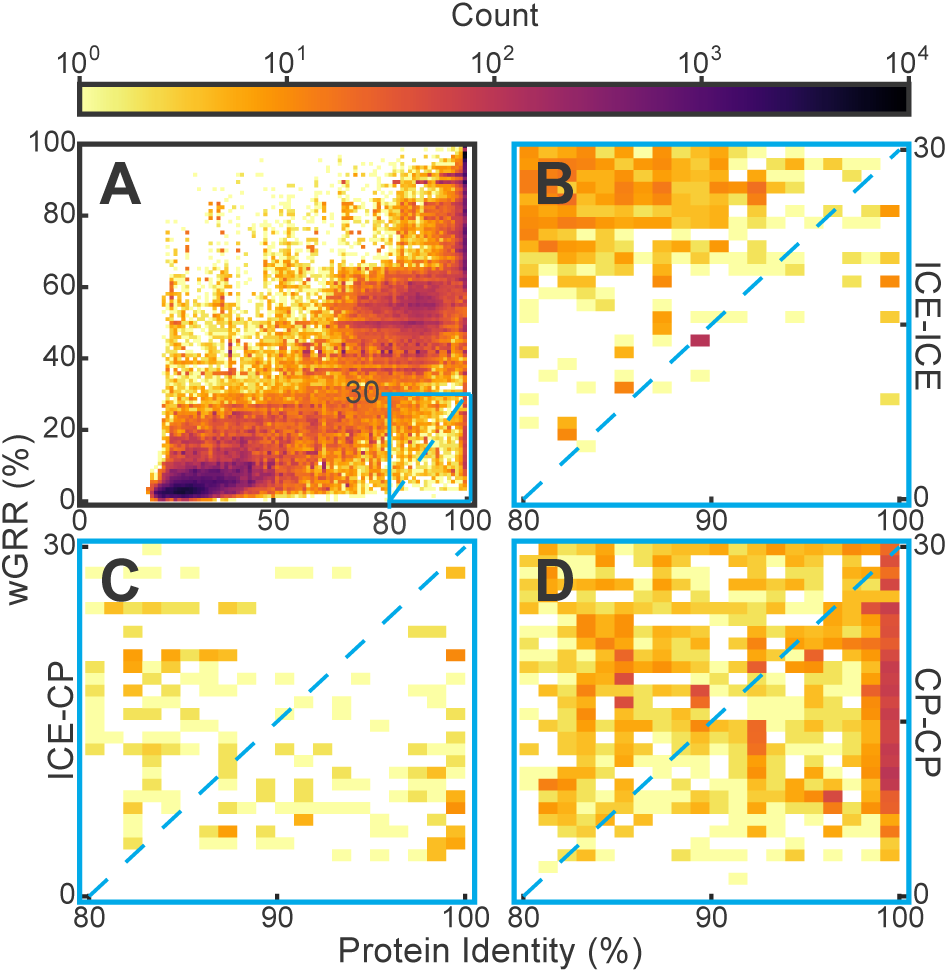
2D histogram of the wGRR score of pairs of conjugative elements as a function of the protein identity of their homologues. **A**. Distribution for the entire dataset. wGRR values are correlated with protein identity of the elements ‘ homologs (ρ _ICE-ICE_ = 0.90, *ρ*_CP-CP_ = 0.83). The blue rectangle zooms on a region where the pairs of elements are very different (GRR<30%), yet they encode at least one very similar protein (identity > 80%). The dashed line separates the elements where protein identity is higher than wGRR × 2/3. **B**. Zoom for ICE-ICE comparisons. **C**. ICE-CP comparisons. **D**. CP-CP comparisons.

We hypothesized that ICEs could hold an advantage over CPs to colonize novel hosts, because replication restricts plasmid host range. To test this hypothesis, we analyzed the wGRR between pairs of ICEs and pairs of CPs in function of the phylogenetic distance between their bacterial hosts. This showed similar patterns for the two types of elements, with the notable exception that there are no pairs of highly similar plasmids (wGRR>50%) in distant hosts (more than 0.1 substitutions/position, *e.g.*, the average distance in the tree between *Escherichia coli* and *Pseudomonas aeruginosa*). In contrast, a third of all ICEs (n=50) are in these conditions (Fig. 6A, Fig. S5). The same analysis after removing the MPF genes shows wGRR values shifted to lower wGRR values for all elements, but qualitatively similar trends (Fig. S6). This suggests a major difference in the ability of ICEs and CPs to be stably maintained after their transfer into a distant host.

**Fig. 6:**
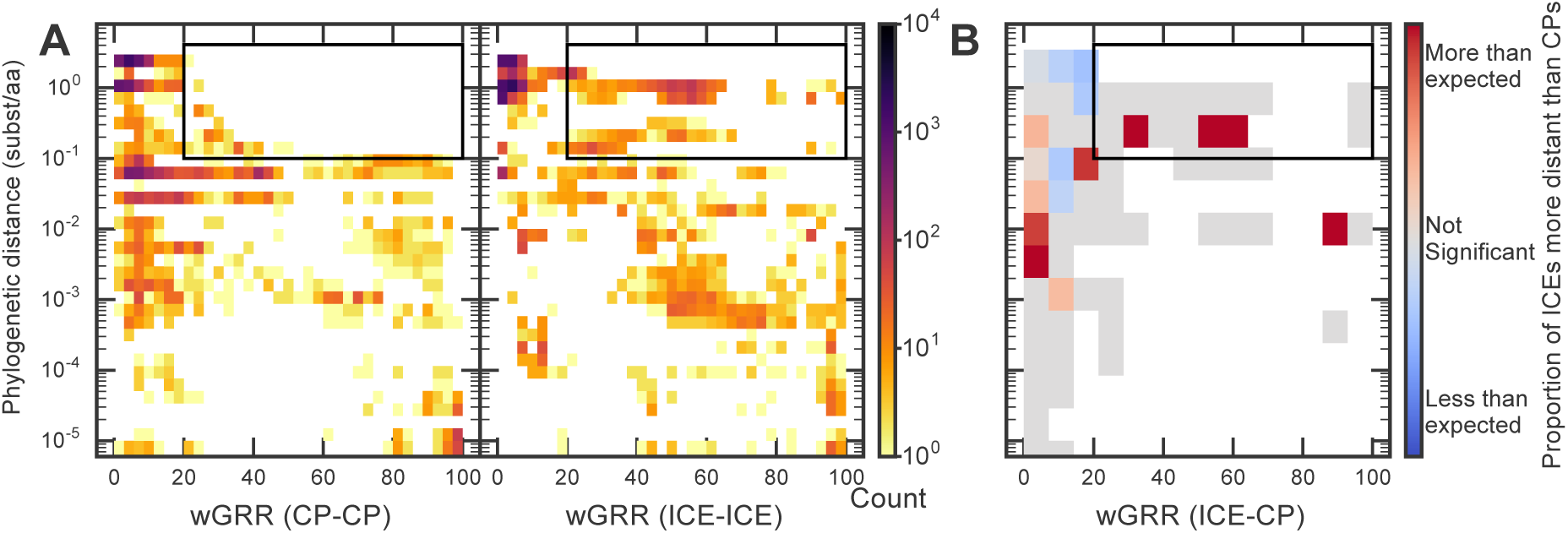
wGRR as function of the host phylogenetic distance **A**. 2D histogram of the distribution of the wGRR score as a function of the phylogenetic distance for pairs of CPs (left) and pairs of ICEs (right). The bottom row corresponds to all pairs with distance lower than 10^−5^ (including those in the same host). Elements in the black rectangle are depicted in a phylogenetic tree in Fig. S5. **B**. Proportion of ICEs more distinct in terms of tri-nucleotides from their host than CPs. Bins are larger than in A. to increase the power of the statistical analysis. Color code: ICEs (Red) or CPs (Blue) are more distinct from the host in terms of tri-nucleotide composition than the other element. Grey: not significant (binomial test, p-value > 10^−2^). White: no elements in the bin.

We then analyzed the pairs ICE-CP. We found few pairs of highly similar ICEs and CPs in closely related hosts (bottom right corner of Fig. 6, n=8 for wGRR>50% and d < 10^−2^), suggesting that interconversion between these elements remains rare within a clade. A larger number of ICE-CP pairs were very similar but present in distant hosts (n=38, Fig. 6). The most parsimonious explanation for these observations, is the recent transfer of one of the elements (ICE or CP) to a distant bacterial host. We identified the latter element based on the differences in terms of tri-nucleotide composition between the elements and the host chromosomes (defined as pvalue in (31)). We then computed for each ICE-CP pair the difference between the pvalue of the pair ICE-host and that of the pair CP-host (see Methods). In agreement to our observation that ICEs have broader host ranges, these differences indicate that ICEs are relatively more distant from the host chromosome for pairs with high wGRR in distant hosts than for the rest of the pairs (Wilcoxon rank sum test, p-value<10^−20^, Fig. 6B, and Fig. S7). The rarity of ICE-CP pairs in closely related hosts, their abundance in distant hosts, and the identification that ICEs are the most compositionally atypical relative to the host in the latter, suggest that successful transfer of CPs to distant hosts is favored when they integrate the chromosome and become ICEs.

## Discussion

In this study, we compared the genetic organization of ICEs and CPs to evaluate the hypothesis that they are essentially equivalent MGEs (15, 16, 29). We found that numerous CPs have integrases (although these may serve for dimer resolution and not integration (32)), and numerous ICEs encode replication and partition functions. Relaxases – present in both ICEs and CPs – have been shown to act as integrases or replication initiators in certain elements (33), which provides further functional overlapping between the elements. Hence, ICEs and CPs share many functions beyond those related to conjugation. These similarities explain why they can cluster together in terms of gene repertoires, even when excluding conjugation functions.

There are also some clear differences between CPs and ICEs explaining why they can also group separately. First, genes encoding replicases and partition systems are more frequent in plasmids, while tyrosine and serine recombinases are more frequent in ICEs. Interestingly, we could not attribute incompatibility groups to ICEs, suggesting that, either the replication module is rarely exchanged between ICEs and CPs, or it evolves too rapidly. Second, the frequency of certain accessory traits is different: plasmids are more likely to encode antibiotic resistance genes whereas ICEs encode more metabolism-related genes. We controlled for taxonomical bias by analyzing ICEs and CPs from the same genera, yet the precise functions we observed are dependent on the dataset which, here, is biased toward nosocomial pathogens. A dataset of bacteria from other environments might present other functional differences in terms of the traits they carry. Finally, the difference in %GC content relative to the host, the number of repeats, the patterns of gene variation and exchange, and the host range are quantitatively different in the two types of elements. After integrating all this information, we propose that, in spite of their similarities, each type has advantages favored in specific situations. In particular, our results suggest that interconversion between ICEs and CPs gives them access to the higher genetic plasticity of CPs and the broader host range of ICEs (Fig. 7).

**Fig. 7:**
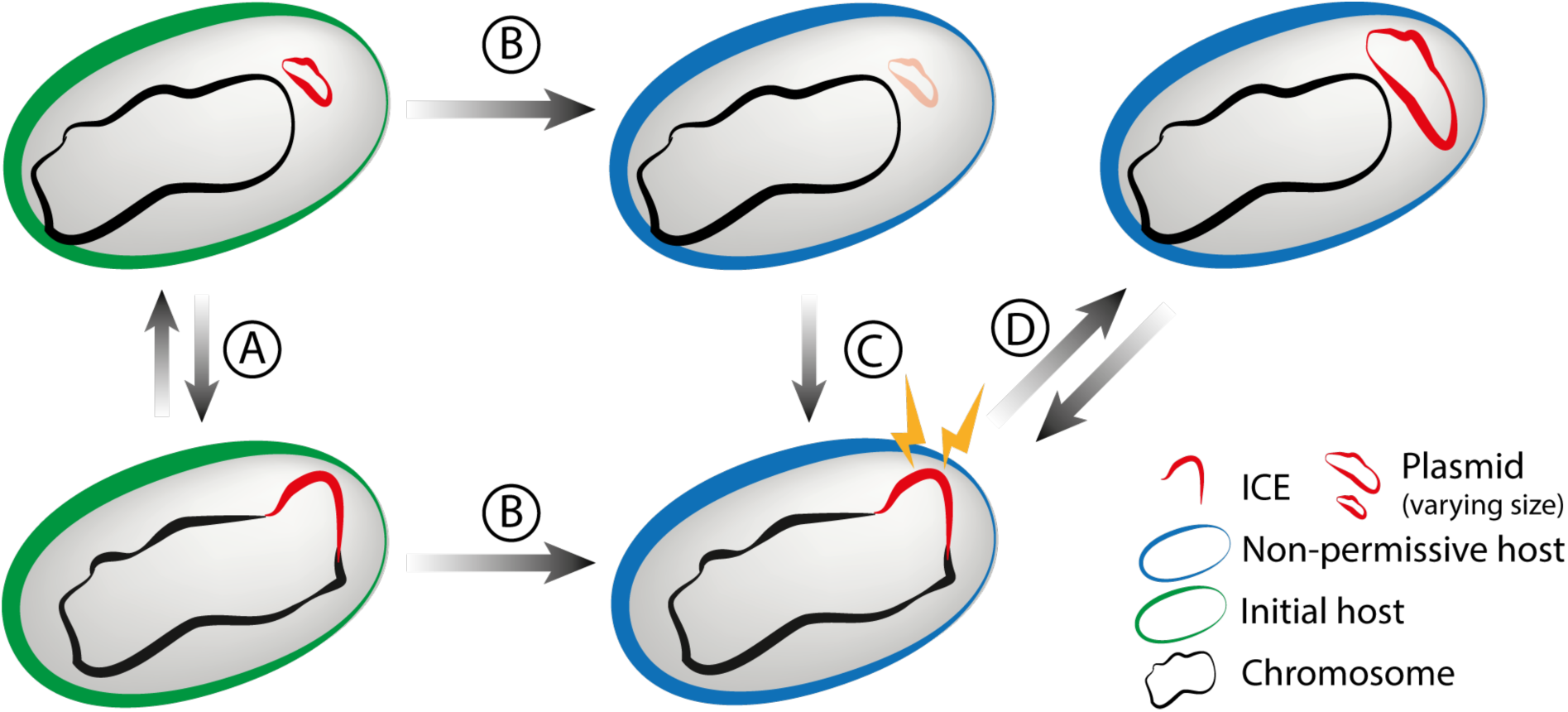
Interconversion between ICEs and CPs allows access to the higher genetic plasticity of CPs and the broader host range of ICEs. **A**. Many ICEs and CPs encode the functions of the other element, facilitating CP<->ICE interconversion. **B**. Conjugative transfer to a taxonomically distant host often precludes stabilization of CPs because they are non-permissive for plasmid replication. ICEs seem at an advantage when conjugating to distant hosts, presumably because they integrate the chromosome. **C**. CPs can be salvaged after transfer in a non-permissive host if they integrate the chromosome (as ICE). We call this effect host-range expansion. **D**. Mutations and/or gene acquisitions can result in an ICE with the ability to replicate as a plasmid. CPs have wider distribution of size and exchange more genetic information, which allows them to acquire novel genetic information at higher rates, which may eventually be adaptive for their hosts.

Even if there are some known families of large (>200kb) ICEs (34, 35), these elements have a remarkably narrower variation in size than CPs in our dataset. This suggests that CPs are more flexible than ICEs in terms of the amount of genetic information they can carry and in their ability to accommodate novel information. We also show that CPs exchange genes more frequently. Mechanistically, this higher rate of gene exchange between CPs may result from recombination between DNA repeats, gene acquisition by integrons, and genome rearrangements by transposable elements. We showed that CPs encode all of these in higher number. Plasmid copy number, when high, may also contribute to increase recombination rates in plasmids relative to ICEs. Interestingly, recombination mediated by transposable elements has been shown to drive the evolution of certain plasmids (36, 37) and to accelerate the reduction of plasmid cost, thus stabilizing the element after horizontal transfer (38). The restrictions in size variation of ICEs are probably not due to the mechanism of integration or excision, because such reactions occur between very distant recombination sites (39). Instead, very large ICEs may disrupt chromosome organization by affecting the distribution of motifs, changing chromosome folding domains, or unbalancing the sizes of replichores (40). Repeat-mediated recombination leads to replicon rearrangements, and may lead to stronger counter-selection of DNA repeats in ICEs than in CPs, further restricting their size variation and their flux of gene exchange. Interestingly, the size of plasmids varies much more steeply with genome size than the size of ICEs, suggesting that CPs may play a particularly important role in the evolution of large bacterial genomes of Proteobacteria, which have higher rates of genetic exchanges (41), and often contain mega-plasmids (42).

Some plasmids are known to be broad-host range and adapt to novel hosts, especially if they carry adaptive traits that compensate for the initially poor intrinsic persistence of the element (43, 44). However, within the large phylogenetic span considered in this work, MPF_T_ ICEs have broader host ranges than CPs. Actually, the first known ICE, Tn916 (not MPF_T_, thus not included in this study), became notorious due to its ability to spread antibiotic resistance between distant phyla (45). We propose that the plasmid to ICE conversion can elicit conjugative transmission to otherwise non-permissive hosts. This results in an effective expansion of the host range of the element (now an ICE). Once installed in the recipient chromosome, the ICE can incorporate a new replication system, or freely mutate its own. If it obtains a functional replication system in the new host the reverse interconversion may occur (Fig. 7). This results in a plasmid with a new host range and the higher evolvability of an autonomous plasmid.

The similarities between certain groups of ICEs and CPs in terms of gene repertoires, the integration of CPs as ICEs upon long-range transfer, and the exchange of genetic information between them are novel evidence for interconversion between the two types of elements. This had previously been proposed based on the phylogeny of the conjugative system (7). Inter-conversion of elements and occasional transfers between CPs and ICEs allow them to access the other elements’ gene pool. These events may result in conjugative elements sharing many traits of the other type of element, as we observed in more than a third of all conjugative elements, and should produce many genetic similarities between ICEs and CPs. The latter probably facilitate further interconversions between the elements.

Other traits may provide advantages specifically to either ICEs or CPs. The ability of plasmids to modify their copy number may accelerate adaptive evolutionary processes, such as the acquisition of antibiotic resistance (46). On the other hand, ICEs might be more stably maintained in lineages because they replicate within the chromosome. Finally, the carriage of ICEs and CPs may have different costs. The cost of plasmids has been extensively studied and is strongly dependent on the traits they encode (47). Much less is known about the cost of ICEs; several reports suggest that they lead to low fitness costs when conjugation is not expressed, but their fitness cost varies much more between elements during transfer (14). Direct comparisons of the cost of carriage of ICEs and CPs carrying similar traits are unavailable. Further experimental work will be needed to test these hypotheses.

Many mobile elements are mobilizable but not able to conjugate independently (42, 48). These elements often encode a relaxase that recognizes the element‘s origin of transfer and is able to interact with a T4SS from an autonomously conjugative element to transfer to other cells. Some other elements only contain an origin of transfer that is recognized by a relaxase of another element. Many of the disadvantages of CPs and ICEs are similar to those of mobilizable plasmids and integrative mobilizable elements, whether they encode a relaxase or not. Notably, the former must be replicated in the extrachromosomal state, and the latter integrate the genome where they must not disrupt genome organization. Patterns observed in conjugative elements are thus likely to be applicable to mobilizable ones.

These results may also be relevant to understand lysogeny by temperate phages. The vast majority of known prophages are integrated in the chromosome, but some replicate like plasmids (49, 50). Considering that prophages share some of the constraints of conjugative elements, they are likely to be under similar trade-offs. However, phages are under additional constraints. Notably, their genome size is much less variable than that of conjugative elements, because it must be packaged into the virion (51), and this may render the extrachromosomal prophages less advantageous in terms of accumulating novel genes. This could explain why most prophages are integrative whereas conjugative systems are more evenly split between integrative and extrachromosomal elements.

In summary, our results show that there are specific fitness benefits associated with the divergent lifestyles identified for pairs of highly similar ICEs and CPs. We should emphasize that our model proposes that plasmid to ICE transition results in broadening the host range of the element, with a concomitant fitness benefit associated with higher rates of its horizontal transfer. The ICE to plasmid transition, on the other hand, results in added versatility of the cargo content (increased size range, etc.), which enhances the evolvability of the mobile element. These factors may promote or even drive the interconversion between plasmids and ICEs in an ever-changing environment. The concepts presented in this work will provide a better understanding of the evolution of bacterial genomes and their mobile genetic elements.

## Material and Methods

### Data

Conjugative systems of type T (MPF_T_) were searched in the set of complete bacterial genomes from NCBI RefSeq (http://ftp.ncbi.nih.gov/genomes/refseq/bacteria/, last accessed in November 2016). We analyzed 5562 complete genomes from 2268 species, including 4345 plasmids and 6001 chromosomes. The classification of the replicon in plasmid or chromosome was taken from the information available in the GenBank file. Our method to delimit ICEs is based on comparative genomics of closely related strains. Hence, we restricted our search for conjugative systems to the species for which we had at least five genomes completely sequenced (164 species, 2990 genomes).

### Detection of conjugative systems and delimitation of ICEs

Conjugative systems were detected using the CONJscan module of MacSyFinder (52), with protein profiles and definitions of the MPF type T, published previously (53). ICEs were delimited with the same methodology, as developed in a previous work (29). Briefly, we identified the core genomes of the species. The region between two consecutive genes of the core genome defined an interval in each chromosome. We then defined spots as the sets of intervals in the chromosome flanked by genes of the same two families of the core genome (24). We then identified the intervals and the spots with conjugative systems. The information on the sets of gene families of the spots with ICEs (i.e., the spot pan-genome) was used to delimit the element boundaries (script available at https://gitlab.pasteur.fr/gem/spot_ICE). This methodology was shown to be accurate at the gene level (precise nucleotide boundaries are not identifiable by this method, see (29)).

### Functional analyses

Partition systems, replication systems, entry-exclusion systems and restriction modification systems were annotated with HMM profiles, as described in our previous work (29, 54). Integrases were annotated with the PFAM profile PF00589 for the Tyrosine recombinases and the combination of PFAM profiles PF00239 and PF07508 for Serine recombinases. DDE Transposases were detected with Macsyfinder (52) with models used previously (55). Antibiotic resistance genes were detected with ResFams profiles (core version v1.1) (56) using the --cut_ga option. We determined the functional categories of genes using their annotation as provided by their best hits to the protein profiles of the EggNOG database for bacteria (version 4.5, bactNOG) (57). Genes not annotated by the EggNOG profiles were classed as “Unknown” and included in the “Poorly characterized” group. The HMM profiles were used to search the genomes with HMMER 3.1b2 (58), and we retrieved the hits with an e-value lower than 10^−3^ and with alignments covering at least 50% of the profile. Integrons were detected using IntegronFinder version 1.5.1 with the --local_max option for higher accuracy (59). Repeats (direct and inverted) were detected with Repseek (version 6.6) (60) using the option -p 0.001 which set the p-value for determining the minimum seed length.

### Statistics

We tested the over-representation of a given function or group of functions using Fisher‘s exact tests on contingency tables. For partition, replication and integration, the contingency table was made by splitting replicons in those encoding or not encoding the function and between ICEs and CPs. The use of presence/absence data instead of the absolute counts was made because the presence of at least one occurrence of a system is sufficient to have the function and because the counts were always low. For the other functions, the contingency table was made by splitting the proteins of the element in those annotated for a given function and the remaining ones. This allowed to take into account the differences in the number of genes between elements. The Fisher-exact tests were considered as significant after sequential Holm-Bonferroni correction, with a family-wise error rate of 5% (the probability of making at least one false rejection in the multiple tests, the type I error). From the contingency table, we computed the relative ratio (or relative risk) of having a given function more often in ICEs than in CPs. The relative ratio is computed as follow: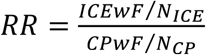where *ICEwF* is the number of ICE (or proteins in ICEs) with the given function, and *N*_*ICE*_, the total number of ICE (or proteins in ICEs), and likewise for CP. The term *ICEwF/N*_*ICE*_is an estimation of the probability of an ICE (or a protein in an ICE) to carry a given a function.

### Phylogenetic distances

Phylogenetic distances were extract from the Proteobacterial tree of the Core-genome. To build the tree, we identified the genes present in at least 90% of the 2897 genomes of Proteobacteria larger than 1 Mb that were available in GenBank RefSeq in November 2016. A list of orthologs was identified as reciprocal best hits using end-gap free global alignment. Hits with less than 37% similarity in amino acid sequence and more than 20% difference in protein length were discarded. We then identified the protein families with relations of orthology in at least 90% of the genomes. They represent 341 protein families. We made multiple alignments of each protein family with MAFFT v.7.205 (with default options) (61) and removed poorly aligned regions with BMGE (with default options) (62). Genes missing in a genome were replaced by stretches of “-” in each multiple alignment, which has been shown to have little impact in phylogeny reconstruction (63). The tree of the concatenate alignment was computed with FastTree version 2.1 under the LG model (64). We chose the LG model because it was the one that minimized the AIC.

### Distance to the host

We used the differences in tri-nucleotide composition to compute the genetic distance between the mobile element and its host chromosome, as previously proposed (65). The analysis of ICEs was done by comparing the element with the chromosome after the removal of its sequence from the latter. Briefly, we computed the trinucleotide relative abundance *χ_ijk_∀i,j,k ∈*{*A,T,C,G*}) for the chromosomes (in windows of 5 kb) and for the conjugative elements (entire replicon), which is given by: *χ*_*ijk*_*=f*_*ijk*_*/f*_*i*_*f*_*i*_*f*_*i*_, with *f* the frequency of a given k-mer in the sequence (31). We first computed the Mahalanobis distance between each window and the host chromosome as follow:

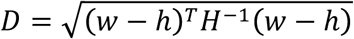

with *w*, the vector of tri-nucleotide abundances (*χ*_*ijk*_) in a given window, and *h*, the mean of the vector of *χ_ijk_* (*i.e.*, the average tri-nucleotide abundance in the chromosome). *H* is the covariance matrix of the tri-nucleotide relative abundances. The inverse of the covariance matrix (H^-1^) downweights frequent trinucleotides, like the tri-nucleotides corresponding to start codons, which are common to conjugative elements and chromosome and could bias the distance. We computed the Mahalanobis distance between conjugative elements and their hosts’ chromosomes (same formula as above, but *w* is now for a conjugative element instead of a chromosome window). We then computed the probability (p-value) that the measured distance between a conjugative element and the host‘s chromosome is the same as any fragment of the host‘s chromosome.

We compared ICEs and CPs in relation to their compositional distance to the host. For this, we made the null hypothesis that the proportion of ICEs having a p-value lower than CPs follows a binomial distribution whose expected proportion is that of the entire dataset (the proportion of ICEs having a p-value lower than CP), precisely: *H*_0_ = *N*(*pvalue_ICE_* < *pvalue_ICE_*)/*N_Comparisons_*, where *N_Comparisons_* is the total number of ICE-CP pairs, *i.e.* 151×136 = 20536.

### Network analysis of gene repertoire relatedness

We built a network describing the relations of homology between the elements. The nodes in the network are conjugative elements and they are linked if they share a given relation of homology. More precisely, the relationship between two elements was quantified with the weighted Gene Repertoire Relatedness score (wGRR). This score represents the number of homologous proteins between two elements, weighted by their sequence identity, as described in (29). The formula is:

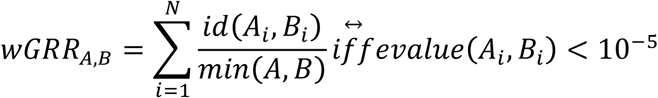

Where (*A*_*i*_, *B*_*i*_) is the *i^th^* pair among N pairs of homologous proteins between element A and element B, id(A_i_, B_i_) is the sequence identity of their alignment, min(A,B) is the number of proteins of the element with fewest proteins (A or B). The sequence identity was computed with blastp v.2.2.15 (default parameters)(66) and kept all bi-directional best hits with an e-value lower than 10^−5^.

The network was built based on the wGRR matrix. Its representation was made using the Fruchterman-Reingold force-directed algorithm as implemented in the NetworkX v1.11 python library. The groups were made using the Louvain algorithm (30). We controlled for the consistency of the heuristic used to assess that the group found are not form a local optimal. We performed 100 clustering, which led to the same classification in 95% of the time.

### Incompatibility typing

We determined the incompatibility group of replicons using the method of PlasmidFinder (67). We used BLASTN (66) to search the replicons for sequences matching the set of 116 probes used by PlasmidFinder. We kept the hits with a coverage above 60% and sequence identity above 80%, as recommended by the authors. Around 3% of the elements had multiple incompatibility types attributed.

## Acknowledgements

This work was supported by an European Research Council grant [EVOMOBILOME n°281605], and a grant from the Agence National de la Recherche [MAGISBAC, ANR-14- CE10-0007]. Work in FdlC lab was supported by grants BFU2014-55534-C2-1-P and BFU2014-62190-EXP from the Spanish Ministry of Economy and Competitiveness. We thank Alan Grossman and Marie Touchon for comments and suggestions, and Aude Bernheim for providing the phylogenetic tree of proteobacteria. J.C. is a member of the ‘Ecole Doctorale Frontière du Vivant (FdV) – Programme Bettencourt‘.

